# Robust effects of working memory demand during naturalistic language comprehension in language-selective cortex

**DOI:** 10.1101/2021.09.18.460917

**Authors:** Cory Shain, Idan A. Blank, Evelina Fedorenko, Edward Gibson, William Schuler

## Abstract

A standard view of human language processing is that comprehenders build richly structured mental representations of natural language utterances, word by word, using computationally costly memory operations supported by domain-general working memory resources. However, three core claims of this view have been questioned, with some prior work arguing that (1) rich word-by-word structure building is not a core function of the language comprehension system, (2) apparent working memory costs are underlyingly driven by word predictability (*surprisal*), and/or (3) language comprehension relies primarily on domain-general rather than domain-specific working memory resources. In this work, we simultaneously evaluate all three of these claims using naturalistic comprehension in fMRI. In each participant, we functionally localize (a) a language-selective network and (b) a ‘multiple-demand’ network that supports working memory across domains, and we analyze the responses in these two networks of interest during naturalistic story listening with respect to a range of theory-driven predictors of working memory demand under rigorous surprisal controls. Results show robust surprisal-independent effects of word-by-word memory demand in the language network and no effect of working memory demand in the multiple demand network. Our findings thus support the view that language comprehension (1) entails word-by-word structure building using (2) computationally intensive memory operations that are not explained by surprisal. However, these results challenge (3) the domain-generality of the resources that support these operations, instead indicating that working memory operations for language comprehension are carried out by the same neural resources that store linguistic knowledge.

**Significance Statement:** This study uses fMRI to investigate signatures of working memory (WM) demand during naturalistic story listening, using a broad range of theoretically motivated estimates of WM demand. Results support a strong effect of WM demand in language-selective brain regions but no effect of WM demand in “multiple demand” regions that have previously been associated with WM in non-linguistic domains. We further show evidence that WM effects in language regions are distinct from effects of word predictability. Our findings support a core role for WM in incremental language processing, using WM resources that are specialized for language.

## Introduction

What computations allow humans to build mental representations of the sentences they are processing, word by word (e.g., Tanenhaus et al., 1995)? A standard view is that richly structured linguistic representations are incrementally stored, retrieved, and integrated in working memory (WM) via computationally intensive operations (e.g., Frazier & Fodor, 1978; Clifton & Frazier, 1989; Just & Carpenter, 1992; Gibson, 2000; Lewis & Vasishth, 2005; Pallier et al., 2011; see **Figure 1**). The WM resources that support these operations are often regarded as domain general, either implicitly by invocation of general constraints on WM (e.g., Gibson, 2000; van Schijndel et al., 2013) or explicitly based on experimental results (e.g., Gordon et al., 2002; Fedorenko et al., 2006, 2007). This domain-general construal of WM for language is consistent with neural mechanisms for abstract structure building in language and other domains of cognition (e.g., Just & Carpenter, 1992; Patel, 2003; Novick et al., 2005).

**Figure 1:**
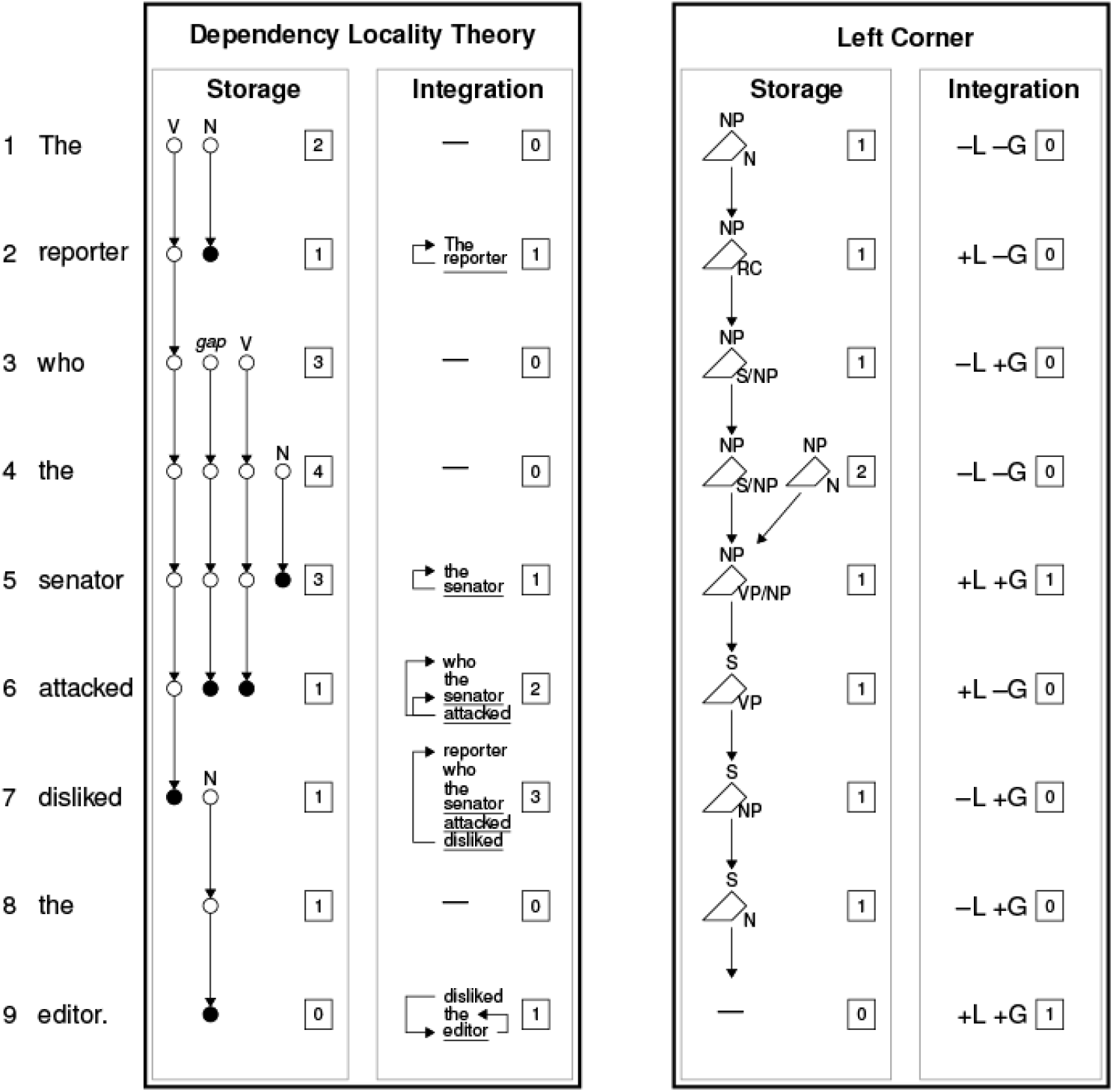
Visualization of storage and integration and their associated costs in two of the three frameworks investigated here: the Dependency Locality Theory (DLT; Gibson, 2000) vs. left corner parsing theory (e.g., Rasmussen & Schuler, 2018). (The third framework—ACT-R (Lewis & Vasishth, 2005)—assumes a left corner parsing algorithm as in the figure above but differs in predicted processing costs, positing (a) no storage costs and (b) integration costs continuously weighted both by the recency of activation for the retrieval target and by the degree of retrieval interference.) Costs are shown in boxes at each step. **DLT walk-through**. In the DLT, expected incomplete dependencies (open circles) are kept in WM and incur storage costs (SC), whereas dependency construction (closed circles) requires retrieval from WM of the previously encountered item and incurs integration costs (IC). Discourse referents (DRs; effectively, nouns and verbs) that contribute to integration costs are underlined in the figure. At *The*, the processor hypothesizes and stores both an upcoming main verb for the sentence (V) and an upcoming noun complement (N). At *reporter*, the expected noun is encountered, contributing 1 DR and a dependency from *reporter* to *the*, which frees up memory. At *who*, the processor posits both a relative clause verb and a gap site, which is coreferent with *who*, and an additional noun complement is posited at *the*. The expected noun is observed at *senator*, contributing 1 DR and a dependency from *senator* to *the*. The awaited verb is observed at *attacked*, contributing 1 DR and two dependencies, one from *attacked* to *senator* and one from the implicit object gap to *who*. The latter spans 1 DR, increasing IC by 1. When *disliked* is encountered, an expected direct object is added to storage, and a subject dependency to *reporter* is constructed with an IC of 3 (the DR *disliked*, plus 2 intervening DRs). At the awaited object *editor*, the store is cleared and two dependencies are constructed (to *the* and *disliked*). **Left corner walk-through**. The memory store contains one or more incomplete derivation fragments (shown as polygons), each with an active sign (top) and an awaited sign (right) needed to complete the derivation. Storage cost is the number of derivation fragments currently in memory. Integration costs derive from binary lexical match (L) and grammatical match (G) decisions. Costs shown here are *end of multiword center embedding* (+L+G), where disjoint fragments are unified (though other cost definitions are possible, see below). At *the*, the processor posits a noun phrase (NP) awaiting a noun. There is nothing on the store so both match decisions are negative. At *reporter*, the noun is encountered (+L) but the sentence is not complete (-G), and the active and awaited signs are updated to NP and relative clause (RC), respectively. At *who*, the processor updates its awaited category to S/NP (sentence with gapped/relativized NP). When *the* is encountered, it is analyzed neither as S/NP nor as a left child of a S/NP; thus both match decisions are negative and a new derivation fragment is created in memory with active sign NP and awaited sign N. Lexical and grammatical matches occur at *senator*, unifying the two fragments in memory, and the awaited sign is updated to VP/NP (VP with gapped NP, the missing unit of the RC). The awaited VP (with gapped NP) is found at *attacked*, leading to a lexical match, and the awaited sign is updated to the missing VP of the main clause. The next two words (*disliked* and *the*) can be incorporated into the existing fragment, updating the awaited sign each time, and *editor* satisfies the awaited N, terminating the parse. **Comparison**. Both approaches posit storage and integration (retrieval) mechanisms but they differ in the details. For example, the DLT (but not left corner theory) posits a spike in integration cost at *attacked*. Differences in predictions between the two frameworks fall out from different claims about the role of WM in parsing.

However, critical tenets of this standard view have been challenged by prior work. In particular, (1) comprehenders may not typically perform rich word-by-word structure building (Swets et al., 2008; Frank & Bod, 2011; Frank & Christiansen, 2018); (2) even if they do, the dominant cost may be due to unexpected input (*surprisal*, a measure of word predictability), not WM demand (e.g., Hale, 2001; Levy, 2008); and (3) even if WM demand is a core determinant of comprehension difficulty, the WM resources may not be the same as those used in non-linguistic tasks like serial recall (e.g., Lewis, 1996; Caplan & Waters, 1999; Fiebach et al., 2001; Santi & Grodzinsky, 2007).

In order to argue that domain-general WM plays a central role in human language comprehension, it is therefore critical both to control for surprisal as an alternative explanation and to determine whether the brain regions that support WM across domains also register WM effects during language comprehension. Therefore, here we use data from a previous large-scale naturalistic fMRI study (Shain, Blank, et al., 2020) to explore multiple existing theories of WM in language processing, under rigorous surprisal controls (van Schijndel & Linzen, 2018). We then evaluate the most robust of these on unseen data. To evaluate the nature of the WM resources that support language comprehension, we examine two candidate brain networks, each functionally localized in individual participants: the language-selective network (Fedorenko et al., 2011) and the domain-general multiple-demand (MD) network implicated in executive functions, including working memory (Duncan et al., 2020).

To foreshadow the key results, responses in the MD network show no evidence of an association with any of the estimators of working memory load explored here. In contrast, responses in the language network show clear and generalizable WM effects, especially as described by the Dependency Locality Theory (Gibson, 2000), whereby the cost of retrieving previously encountered dependents of a currently processed word is determined by the amount of intervening material. Based on these results, we argue (a) that a core function of the human language processor is to compose representations in working memory based on structural cues, even for naturalistic materials during passive listening, and (b) that these operations are implemented locally within the language-selective cortical network. Our study thus supports the view that typical language comprehension involves rich word-by-word structure building via computationally intensive memory operations but challenges the domain-generality of the neural resources that implement these memory operations, in line with recent arguments against a separation between storage and computation in the brain (e.g., Hasson et al., 2015; Dasgupta & Gershman, 2021).

## Materials and Methods

Except where otherwise noted below, we use the materials and methods of Shain, Blank, et al. (2020). At a high level, we analyze the influence of theory-driven measures of working memory load during auditory comprehension of naturalistic stories (Futrell et al., 2020) on activation levels in the language-selective (LANG) vs. domain-general multiple-demand (MD) networks identified in each participant using an independent functional localizer. To control for regional and participant-level variation in the hemodynamic response function (HRF), the HRF is estimated from data using continuous-time deconvolutional regression (Shain & Schuler, 2018, 2021), rather than assumed (cf. e.g., Bhattasali et al., 2019; Brennan et al., 2016). Hypotheses are tested using generalization performance on held-out data.

### Experimental Design

Functional magnetic resonance imaging data were collected from seventy-eight native English speakers (30 males), aged 18-60 (*M*±*SD* = 25.8±9, *Med*±*SIQR* = 23±3). Each participant completed a passive story comprehension task, using materials from Futrell et al. (2020), and a functional localizer task designed to identify the language and MD networks and ensure functionally comparable units of analysis across participants. The use of functional localization is motivated by established inter-individual variability in the precise locations of functional areas (e.g., Frost & Goebel, 2012; Tahmasebi et al., 2012; Vázquez-Rodríguez et al., 2019)—including the LANG (e.g., Fedorenko et al., 2010) and MD (e.g., Fedorenko et al., 2013; Shashidhara et al., 2020) networks. Functional localization yields higher sensitivity and functional resolution compared to the traditional voxel-wise group-averaging fMRI approach (e.g., Nieto-Castañón & Fedorenko, 2012), and is especially important given the proximity of the LANG and the MD networks in the left frontal cortex (see Fedorenko & Blank, 2020, for a review). The localizer task contrasted sentences with perceptually similar controls (lists of pronounceable non-words). Participant-specific functional regions of interest (fROIs) were identified by selecting the top 10% of voxels that were most responsive (to the target contrast; see below) for each participant within broad areal ‘masks’ (derived from probabilistic atlases for the same contrasts, created from large numbers of individuals; see Fedorenko et al., 2010, for the description of the general approach).

Six left-hemisphere language fROIs were identified using the contrast *sentences > nonwords*: in the inferior frontal gyrus (IFG) and its orbital part (IFGorb), in the middle frontal gyrus (MFG), in the anterior and posterior temporal cortex (AntTemp and PostTemp), and in the angular gyrus (AngG). This contrast targets higher-level aspects of language, to the exclusion of perceptual (speech / reading) and motor-articulatory processes (for discussion, see Fedorenko & Thompson-Schill, 2014, or Fedorenko, 2020). Critically, this localizer has been extensively validated over the past decade across diverse parameters and shown to generalize across task (Fedorenko et al., 2010; Ivanova et al., 2020), presentation modality (Fedorenko et al., 2010; Scott et al., 2017; Chen, Affourtit et al., 2021), language (Ayyash, Malik-Moraleda et al., 2021), and materials (Fedorenko et al., 2010; Ivanova et al., 2020), including both coarser contrasts (e.g., between natural speech and an acoustically degraded control: Scott et al., 2017) and narrower contrasts (e.g., between lists of unconnected, real words and nonwords lists, or between sentences and lists of words: Blank et al., 2016; Fedorenko et al., 2010). Moreover, a network that corresponds to the one identified by the *sentences > nonwords* contrast emerges robustly from task-free (resting state) data using data-driven functional correlation approaches (Braga et al., 2020; see also Blank et al., 2014), suggesting that this network is a stable ‘natural kind’ and the use of functional localizers is simply a quick and efficient way to identify this network.

Ten multiple-demand fROIs were identified bilaterally using the contrast *nonwords > sentences*, which reliably localizes the MD network, as discussed below: in the posterior (PostPar), middle (MidPar), and anterior (AntPar) parietal cortex, in the precentral gyrus (PrecG), in the superior frontal gyrus (SFG), in the middle frontal gyrus (MFG) and its orbital part (MFGorb), in the opercular part of the inferior frontal gyrus (IFGop), in the anterior cingulate cortex and pre-supplementary motor cortex (ACC/pSMA), and in the insula (Insula). This contrast targets regions that increase their response with the more effortful reading of nonwords compared to that of sentences. This “cognitive effort” contrast robustly engages the MD network and can reliably localize it (Fedorenko et al., 2013). Moreover, it generalizes across a wide array of stimuli and tasks, both linguistic and non-linguistic, including, critically, standard contrasts targeting executive functions such as working-memory and inhibitory control (Fedorenko et al., 2013; Mineroff, Blank, et al., 2018; Shashidara et al., 2019). Like the LANG network, the MD network is strongly dissociated from other brain networks during naturalistic cognition (Blank et al., 2014; Paunov et al., 2019; Assem et al., 2020; Braga et al., 2020). To verify that the use of a flipped language localizer contrast does not artificially suppress language-processing-related effects in MD, we performed a follow-up analysis where the MD network was identified with a *hard* > *easy* contrast in a spatial working memory paradigm, which requires participants to keep track of more vs. fewer spatial locations within a grid (e.g., Fedorenko et al., 2013) in the subset of participants (∼80%) who completed this task.

Full details about participants, stimuli, functional localization, data acquisition, and preprocessing are provided in Shain, Blank, et al. (2020).

### Statistical analysis

This study uses continuous-time deconvolutional regression (CDR) for all statistical analyses (Shain & Schuler, 2021). CDR uses machine learning to estimate continuous-time impulse response functions (IRFs) that describe the influence of observing an event (word) on a response (BOLD signal change) as a function of their distance in continuous time. When applied to fMRI, CDR-estimated IRFs represent the hemodynamic response function (Boynton et al., 1996) and can account for regional differences in response shape (Handwerker et al., 2004) directly from responses to naturalistic language stimuli, which are challenging to model using discrete-time techniques (Shain & Schuler, 2021). Model details are as follows.

#### Control Predictors

We include all control predictors used in Shain, Blank, et al. (2020), namely:

- ***Sound Power***: Frame-by-frame root mean squared energy (RMSE) of the audio stimuli computed using the Librosa software library (McFee et al., 2015).
- ***TR Number***: Integer index of the current fMRI volume within the current scan.
- ***Rate***: Deconvolutional intercept. A vector of ones time-aligned with the word onsets of the audio stimuli. *Rate* captures influences of stimulus *timing* independently of stimulus *properties* (see e.g., Brennan et al., 2016; Shain & Schuler, 2018).
- ***Frequency***: Corpus frequency of each word computed using a KenLM unigram model trained on Gigaword 3. For ease of comparison to surprisal, frequency is represented here on a surprisal scale (negative log probability), such that larger values index less frequent words (and thus greater expected processing cost).
- ***Network***: Numeric predictor for network ID, 0 for MD and 1 for LANG. Used only in models of combined responses from both networks.

Furthermore, because points of predicted retrieval cost may partially overlap with prosodic breaks between clauses, we include two prosodic controls:

- ***End of Sentence*:** Indicator for whether a word terminates a sentence.
- ***Pause Duration*:** Length (in ms) of pause following a word, as indicated by hand-corrected word alignments over the auditory stimuli. Words that are not followed by a pause take the value 0ms.

We confirmed empirically that the pattern of significance reported in Shain, Blank, et al. (2020) holds in the presence of these additional controls.

In addition, inspired by evidence that word predictability strongly influences blood oxygen level dependent (BOLD) responses in the language network, we additionally include the critical surprisal predictors from Shain, Blank, et al. (2020):

- ***5-gram Surprisal*:** 5-gram surprisal for each word in the stimulus set from a KenLM (Heafield et al., 2013) language model with default smoothing parameters trained on the Gigaword 3 corpus (Graff et al., 2007). 5-gram surprisal quantifies the predictability of words as the negative log probability of a word given the four words preceding it in context.
- ***PCFG Surprisal*:** Lexicalized probabilistic context-free grammar surprisal computed using the incremental left corner parser of van Schijndel et al. (2013) trained on a generalized categorial grammar (Nguyen et al., 2012) reannotation of Wall Street Journal sections 2 through 21 of the Penn Treebank (Marcus et al., 1993).

PCFG and 5-gram surprisal were investigated by Shain, Blank, et al. (2020) because their interpretable structure permitted testing of hypotheses of interest in that study. However, their strength as language models has now been outstripped by less interpretable but better performing incremental language models based on deep neural networks (Jozefowicz et al., 2016; Gulordava et al., 2018; Radford et al., 2019). In the present investigation, predictability effects are a control rather than an object of study, and we are therefore not bound by the same interpretability considerations. To strengthen the case for independence of retrieval processes from prediction processes, we therefore additionally include the following predictability control:

- ***Adaptive Surprisal*:** Word surprisal as computed by the adaptive recurrent neural network (RNN) of van Schijndel & Linzen (2018). This network is equipped with a cognitively-inspired mechanism that allows it to adjust its expectations to the local discourse context at inference time, rather than relying strictly on knowledge acquired during the training phase. Compared to strong baselines, results show both improved model perplexity and improved fit between model-generated surprisal estimates and measures of human reading times. Because the RNN can in principle learn both (1) the local word co-occurrence patterns exploited by 5-gram models and (2) the structural features exploited by PCFG models, it competes for variance in our regression models with the other surprisal predictors, whose effects are consequently attenuated relative to Shain, Blank, et al. (2020).

Models additionally included the mixed-effects random grouping factors *Participant* and *fROI*. We examine the responses for each network (LANG, MD) as a whole, which is reasonable given strong evidence of functional integration among the regions of each network (e.g., Blank et al., 2014; Braga et al., 2020; Assem et al., 2020), but we also examine each individual fROI separately for a richer characterization of the observed effects. Prior to regression, all predictors were rescaled by their standard deviations in the training set except *Rate* (which has no variance) and the indicators *End of Sentence* and *Network*. Reported effect sizes are therefore in standard units.

#### Critical predictors

Among the many prior theoretical and empirical investigations of working memory demand in sentence comprehension, we have identified three theoretical frameworks that are *broad-coverage* (i.e., sufficiently articulated to predict word-by-word memory demand in arbitrary utterances) and *implemented* (i.e., accompanied by algorithms that can generate word-by-word memory predictors for our naturalistic English-language stimuli): the **Dependency Locality Theory** (DLT; Gibson, 2000), **ACT-R** sentence processing theories (Lewis & Vasishth, 2005), and **left corner parsing** theories (Johnson-Laird, 1983; Resnik, 1992; van Schijndel et al., 2013; Rasmussen & Schuler, 2018). We set aside related work that does not define word-by-word measures of WM demand (e.g., Gordon et al., 2001, 2006; McElree et al., 2003).

At a high level, these theories all posit WM demands driven by the syntactic structure of sentences. In the DLT, the relevant structures are *dependencies* between words (e.g., between a verb and its subject). In ACT-R and left corner theories, the relevant structures are *phrasal hierarchies* of labeled, nested spans of words (syntax trees). The DLT and left corner theories hypothesize active maintenance in memory (and thus ‘storage’ costs) from incomplete dependencies and incomplete phrase structures, respectively, whereas ACT-R posits no storage costs under the assumption that partial derivations live in a content-addressable memory store. All three frameworks posit ‘integration’ costs driven by memory retrieval operations. In the DLT, retrieval is required in order to build dependencies, with cost proportional to the length of the dependency. In ACT-R and left corner theories, retrieval is required in order to unify representations in memory. Left corner theory is compatible with several notions of retrieval cost (explored below), whereas ACT-R assumes retrieval costs are governed by an interaction between continuous-time activation decay mechanisms and similarity-based interference.

Prior work has investigated the empirical predictions of some of these theories using computer simulations (Lewis & Vasishth, 2005; Rasmussen & Schuler, 2018) and human behavioral responses to constructed stimuli (Grodner & Gibson, 2005) and reported robust WM effects. Related work has also shown effects of dependency length manipulations in measures of comprehension and online processing difficulty (e.g., Gibson et al., 1996; McElree et al., 2003; Van Dyke & Lewis, 2003; Makuuchi et al., 2009; Meyer et al., 2013). In light of these findings, evidence from more naturalistic human sentence processing settings for working memory effects of any kind is surprisingly weak. Demberg & Keller (2008) report DLT integration cost effects in the Dundee eye-tracking corpus (Kennedy et al., 2003), but only when the domain of analysis is restricted – overall DLT effects are actually negative (longer dependencies yield shorter reading times, also known as an *anti-locality* effects, Konieczny, 2000). Van Schijndel & Schuler (2013) also report anti-locality effects in Dundee, even under rigorous controls for word predictability phenomena that have been invoked to explain anti-locality effects in other experiments (Konieczny, 2000; Vasishth & Lewis, 2006). It is therefore not yet settled how central syntactically-related working memory involvement is to human sentence processing in general, rather than perhaps being driven by the stimuli and tasks commonly used in experiments designed to test these effects (Hasson & Honey, 2012; Campbell & Tyler, 2018; Hasson et al., 2018; Diachek, Blank, Siegelman et al., 2020). In the fMRI literature, few prior studies of naturalistic sentence processing have investigated syntactic working memory (although some of the syntactic predictors in Brennan et al., 2016, especially syntactic node count, are amenable to a memory-based interpretation).

##### DLT predictors

The DLT posits two distinct sources of WM demand, *integration cost* and *storage cost*. Integration cost is computed as the number of discourse referents that intervene in a backward-looking syntactic dependency, where “discourse referent” is operationalized, for simplicity, as any noun or finite verb. In addition, all the implementation variants of integration cost proposed by Shain et al. (2016) are considered:

- **V:** Verbs are more expensive. Non-finite verbs receive a cost of 1 (instead of 0) and finite verbs receive a cost of 2 (instead of 1).
- **C:** Coordination is less expensive. Dependencies out of coordinate structures skip preceding conjuncts in the calculation of distance, and dependencies with intervening coordinate structures assign that structure a weight equal to that of its heaviest conjunct.
- **M:** Exclude modifier dependencies. Dependencies to preceding modifiers are ignored.

These variants are motivated by the following considerations. First, the reweighting in V is motivated by the possibility (1) that finite verbs may require more information-rich representations than nouns, especially tense and aspect (Binnick, 1991), and (2) that non-finite verbs may still contribute eventualities to the discourse context, albeit with underspecified tense (Lowe, 2019). As in Gibson (2000), the precise weights are unknown, and the weights used here are simply heuristic approximations that instantiate a hypothetical overall pattern: non-finite verbs contribute to retrieval cost, and finite verbs contribute more strongly than other classes.

Second, the discounting of coordinate structures under C is motivated by the possibility that conjuncts are incrementally integrated into a single representation of the overall coordinated phrase, and thus that their constituent nouns and verbs no longer compete as possible retrieval targets. Anecdotally, this possibility is illustrated by the following sentence:

> *Today I bought a cake, streamers, balloons, party hats, candy, and several gifts* ***for*** *my niece’s birthday*.

In this example, the dependency from *for* to its modificand *bought* does not intuitively seem to induce a large processing cost, yet it spans 6 coordinated nouns, yielding an integration cost of 6, which is similar in magnitude to that of some of the most difficult dependencies explored in Grodner & Gibson (2005). The C variant treats the entire coordinated direct object as one discourse referent, yielding an integration cost of 1.

Third, the discounting of preceding modifiers in M is motivated by the possibility that modifier semantics may be integrated early, alleviating the need to retrieve the modifier once the head word is encountered. Anecdotally, this possibility is illustrated by the following sentence:

> *(Yesterday,) my coworker, whose cousin drives a taxi in Chicago*, ***sent*** *me a list of all the best restaurants to try during my upcoming trip*.

The dependency between the verb *sent* and the subject *coworker* spans a finite verb and 3 nouns, yielding an integration cost of 4 (plus a cost of 1 for the discourse referent introduced by *sent*). If the sentence includes the pre-sentential modifier *Yesterday*, which under the syntactic annotation used in this study is also involved in a dependency with the main verb *sent*, then the DLT predicts that it should double the structural integration cost at *sent* because the same set of discourse referents intervenes in two dependencies rather than one. Intuitively, this does not seem to be the case, possibly because the temporal information contributed by *Yesterday* may already be integrated with the incremental semantic representation of the sentence before *sent* is encountered, eliminating the need for an additional retrieval operation at that point. The +M modification instantiates this possibility.

The presence/absence of the three features above yields a total of eight variants: DLT, DLT-V (a version with the V feature), DLT-C, DLT-M, DLT-VC, DLT-VM, DLT-CM, and DLT-VCM. A superficial consequence of the variants with C and M features is that they tend to attenuate large integration costs. Thus, if they improve fit to human measures, it may simply be the case that the DLT in its original formulation overestimates the costs of long dependencies. To account for this possibility, this study additionally considers a log-transformed variant of (otherwise unmodified) DLT integration cost: DLT (log).

We additionally consider DLT storage cost (DLT-S):

- **DLT-S:** The number of awaited syntactic heads at a word that are required to form a grammatical utterance.

In our implementation, this includes dependencies arising via syntactic arguments (e.g., the object of a transitive verb), dependencies from modifiers to following modificands, dependencies from relative pronouns (e.g., *who, what*) to a gap site in the following relative clause, dependencies from conjunctions to following conjuncts, and dependencies from gap sites to following extraposed items. In all such cases, the existence of an obligatory upcoming syntactic head can be inferred from context. This is not the case for the remaining dependency types (e.g., from modifiers to preceding modificands, since the future appearance of a modifier is not required when the modificand is processed), and they are therefore treated as irrelevant to storage cost. Because storage cost does not assume a definition of distance (cf. integration cost), no additional variants of it are explored.

##### ACT-R predictor

The ACT-R model (Lewis & Vasishth, 2005) composes representations in memory through a content-addressable retrieval operation that is subject to similarity-based interference (Gordon et al., 2001; McElree et al., 2003; Van Dyke & Lewis, 2003), with memory representations that decay with time unless reactivated through retrieval. The decay function enforces a locality-like notion (retrievals triggered by long dependencies will on average cue targets that have decayed more), but this effect can be attenuated by intermediate retrievals of the target. Unlike the DLT, ACT-R has no notion of active maintenance in memory (items are simply retrieved as needed) and therefore does not predict a storage cost.

The originally proposed ACTR-R parser (Lewis & Vasishth, 2005) is implemented using hand-crafted rules and deployed on utterances constructed to be consistent with those rules. This implementation does not cover arbitrary sentences of English and cannot therefore be applied to our stimuli without extensive additional engineering of the parsing rules. However, a recently proposed modification to the ACT-R framework has a broad-coverage implementation and has already been applied to model reading time responses to the same set of stories (Dotlačil, 2021). It does so by moving the parsing rules from procedural to declarative memory, allowing the rules themselves to be retrieved and activated in the same manner as parse fragments. In this study, we use the same single ACT-R predictor used in Dotlačil (2021):

- **ACT-R Target Activation:** The mean activation level of the top three most activated retrieval targets cued by a word. Activation decays on both time and degree of similarity with retrieval competitors, and is therefore highest when the cue strongly identifies a recently activated target. *ACT-R target activation* is expected to be anticorrelated with retrieval difficulty. See Dotlačil (2021) and Lewis & Vasishth (2005) for details.

##### Left corner predictors

Another line of research (Johnson-Laird, 1983; Resnik, 1992; van Schijndel et al., 2013; Rasmussen & Schuler, 2018) frames incremental sentence comprehension as left corner parsing (Rosenkrantz & Lewis II, 1970) under a pushdown store implementation of working memory. Under this view, incomplete derivation fragments representing the hypothesized structure of the sentence are assembled word by word, with working memory required to (1) push new derivation fragments to the store, (2) retrieve and compose derivation fragments from the store, and (3) maintain incomplete derivation fragments in the store. For a detailed presentation of a recent instantiation of this framework, see Rasmussen & Schuler (2018). In principle, costs could be associated with any of the parse operations computed by left corner models, as well as (1) with DLT-like notions of storage (maintenance of multiple derivation fragments in the store) and (2) with ACT-R-like notions of retrieval and reactivation, since items in memory (corresponding to specific derivation fragments) are incrementally retrieved and updated. Unlike ACT-R, left corner frameworks do not necessarily enforce activation decay over time, and they do not inherently specify expected processing costs.

Full description of left corner parsing models of sentence comprehension is beyond the scope of this presentation (see e.g., Oh et al., 2021; Rasmussen & Schuler, 2018), which is restricted to the minimum details needed to define the predictors covered here. At a high level, phrasal structure derives from a sequence of lexical match (±L) and grammatical match (±G) decisions made at each word (see Oh et al., 2021 for relations to equivalent terms in the prior parsing literature). In terms of memory structures, the lexical decision depends on whether a new element (representing the current word and its hypothesized part of speech) matches current expectations about the upcoming syntactic category; if so, it is composed with the derivation at the front of the memory store (+L), and if not, it is pushed to the store as a new derivation fragment (-L). Following the lexical decision, the grammatical decision depends on whether the two items at the front of the store can be composed (+G) or not (-G). In terms of phrasal structures, lexical matches index the ends of multiword constituents (+L at the end of a multiword constituent, -L otherwise), and grammatical match decisions index the ends of left-child (center-embedded) constituents (+G at the end of a left child, -G otherwise). These composition operations (+L and +G) instantiate the notion of syntactic “integration” as envisioned by e.g., the DLT, since structures are retrieved from memory and updated by these operations. They each may thus plausibly contribute a memory cost (Shain et al., 2016), leading to the following left corner predictors:

- **End of Constituent (+L):** Indicator for whether a word terminates a multiword constituent (i.e. whether the parser generates a lexical match).
- **End of Center Embedding (+G):** Indicator for whether a word terminates a center embedding (left child) of one or more words (i.e. whether the parser generates a grammatical match).
- **End of Multiword Center Embedding (+L, +G):** Indicator for whether a word terminates a multiword center embedding (i.e. whether the parser generates both a lexical match and a grammatical match).

In addition, the difficulty of retrieval operations could in principle be modulated by locality, possibly due to activation decay and/or interference as argued by Lewis & Vasishth (2005). To account for this possibility, this study also explores distance-based left corner predictors:

- **Length of Constituent (+L):** When a word terminates a multiword constituent, distance from most recent retrieval (including creation) of the derivation fragment at the front of the store (otherwise, 0).
- **Length of Multiword Center Embedding (+L, +G):** When a word terminates a multiword center embedding, distance from most recent retrieval (including creation) of the derivation fragment at the front of the store (otherwise, 0).

The notion of distance must be defined, and three definitions are explored here. One simply counts the number of words (WD). However, this complicates comparison to the DLT, which then differs not only in its conception of memory usage (constructing dependencies vs. retrieving/updating derivations in a pushdown store), but also in its notion of locality (the DLT defines locality in terms of nouns and finite verbs, rather than words). To enable direct comparison, DLT-like distance metrics are also used in the above left corner locality-based predictors – in particular, both using the original DLT definition of discourse referents (DR), as well as the modified variant +V that reweights finite and non-finite verbs (DRV). All three distance variants are explored for both distance-based left corner predictors.

Note that these left corner distance metrics more closely approximate ACT-R retrieval cost than DLT integration cost, because, as stressed by Lewis & Vasishth (2005), decay in ACT-R is determined by the recency with which an item in memory was previously activated, rather than overall dependency length. Left corner predictors can therefore be used to test one of the motivating insights of the ACT-R framework: the influence of reactivation on retrieval difficulty.

Note also that because the parser incrementally constructs expected dependencies between as-yet incomplete syntactic representations, at most two retrievals are cued per word (up to one for each of the lexical and grammatical decisions), no matter how many dependencies the word participates in. This property makes left corner parsing a highly efficient form of incremental processing, a feature that has been argued to support its psychological plausibility (Johnson-Laird, 1983; van Schijndel et al., 2013; Rasmussen & Schuler, 2018).

The aforementioned left corner predictors instantiate a notion of retrieval cost, but the left corner approach additionally supports measures of storage cost. In particular, the number of incomplete derivation fragments that must be held in memory (similar to the number of incomplete dependencies in DLT storage cost) can be read off the store depth of the parser state:

- **Embedding Depth:** The number of incomplete derivation fragments left on the store once a word has been processed.

This study additionally considers the possibility that pushing a new fragment to the store may incur a cost:

- **Start of Embedding (-L-G):** Indicator for whether *embedding depth* increased from one word to the next.

As with retrieval-based predictors, the primary difference between left corner *embedding depth* and DLT *storage cost* is the efficiency with which the memory store is used by the parser. Because expected dependencies between incomplete syntactic derivations are constructed as soon as possible, a word can contribute at most one additional item to be maintained in memory (*vis-a-vis* DLT storage cost, which can in principle increase arbitrarily at words which introduce multiple incomplete dependencies). As mentioned above, ACT-R does not posit storage costs at all, and thus investigation of such costs potentially stands to empirically differentiate ACT-R from DLT/left corner accounts.

#### Model Design

Following Shain, Blank, et al. (2020), we use continuous-time deconvolutional regression (CDR; Shain & Schuler, 2019, 2018) to infer the shape of the hemodynamic response function (HRF) from data (Boynton et al., 1996; Handwerker et al., 2004). We assumed the following two-parameter HRF kernel based on the widely-used double-gamma canonical HRF (Lindquist et al., 2009):

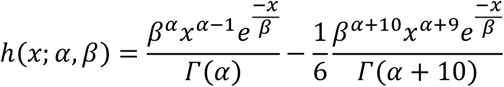

where parameters α, β are fitted using black box variational Bayesian (BBVI) inference. Model implementation follows Shain, Blank, et al. (2020) except in replacing improper uniform priors with normal priors, which have since been shown empirically to produce more reliable estimates of uncertainty (Shain & Schuler, 2021). Variational priors follow Shain & Schuler (2019).

The following CDR model specification was fitted to responses from each of the LANG and MD fROIs, where *italics* indicate predictors convolved using the fitted HRF and **bold** indicates predictors that were ablated for hypothesis tests:

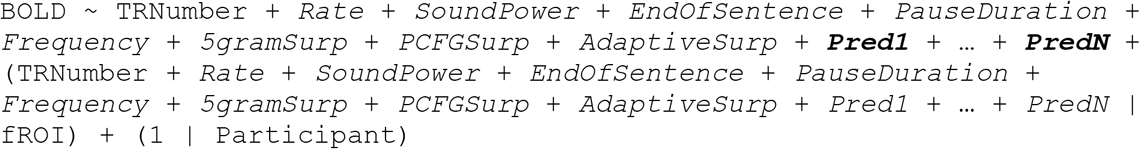

that is, a linear coefficient for the index of the TR in the experiment, convolutions of the remaining predictors with the fitted HRF, by-fROI random variation in effect size and shape, and by-participant random variation in base response level. This model is used to test for significant effects of one or more critical predictors *Pred1*, …, *PredN* in each of LANG and MD. To test for significant *differences* between LANG and MD in the effect size(s) of critical predictor(s) *Pred1*, …, *PredN*, we additionally fitted the following model to the combined responses from both LANG and MD:

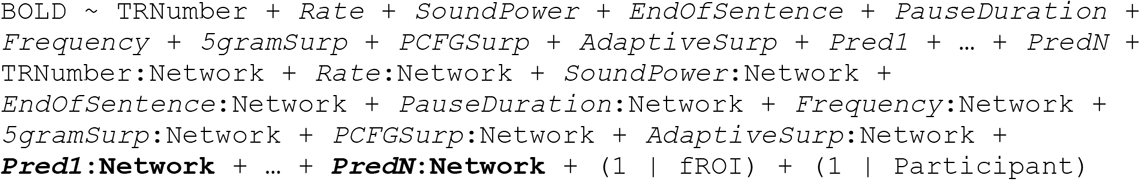

By-fROI random effects are simplified from the individual network models in order to support model identifiability (see Shain, Blank, et al., 2020).

#### Ablative Statistical Testing

Following Shain, Blank, et al. (2020), we partition the fMRI data into training and evaluation sets by cycling TR numbers *e* into different bins of the partition with a different phase for each subject *u*:

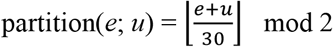

assigning output 0 to the training set and 1 to the evaluation set. Model quality is quantified as the Pearson sample correlation, henceforth *r*, between model predictions on a dataset (training or evaluation) and the true response. Fixed effects are tested by paired permutation test (Demšar, 2006) of the difference in correlation *r*_diff_ = *r*_full_ – *r*_ablated_, where *r*_full_ is the *r* of a model containing the fixed effect of interest while *r*_ablated_ is the *r* of a model lacking it. Paired permutation testing requires an elementwise performance metric that can be permuted between the two models, whereas Pearson correlation is a global metric that applies to the entire prediction-response matrix. To address this, we exploit the fact that the sample correlation can be converted to an elementwise performance statistic as long as both variables are standardized (i.e. have sample mean 0 and sample standard deviation 1):

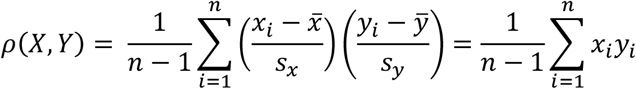

As a result, an elementwise performance metric can be derived as the elements of a Hadamard product between independently standardized prediction and response vectors. These products are then permuted in the usual way, using 10,000 resampling iterations. Each test involves a single ablated fixed effect, retaining all random effects in all models.

#### Exploratory and Generalization Analyses

Exploratory study is needed for the present general question about the nature of WM-dependent operations that support sentence comprehension because multiple broad-coverage theories of WM have been proposed in language processing, as discussed above, each making different predictions and/or compatible with multiple implementation variants, ruling out a single theory-neutral measure of word-by-word WM load. In addition, as discussed in the Introduction, prior naturalistic investigations of WM have yielded mixed results, motivating the use of a broad net to find WM measures that correlate with human processing difficulty. Exploratory analysis can therefore illuminate both the existence and kind of WM operations involved in human language comprehension.

However, exploring a broad space of predictors increases the false positive rate, and thus the likelihood of spurious findings. In order to avoid this issue and enable testing of patterns discovered by exploratory analyses, we divide the analysis into exploratory (in-sample) and generalization (out-of-sample) phases. In the exploratory phase, single ablations are fitted to the training set for each critical variable (that is, a model with a fixed effect for the variable and a model without one) and evaluated via in-sample permutation testing on the training set. This provides a significance test for the contribution of each individual variable to *r*_diff_ in the training set. This metric is used to select models from broad “families” of predictors for generalization-based testing, where the members of each family constitute implementation variants of the same underlying idea:

- DLT integration cost:
  ∘ DLT-(V)(C)(M)
- DLT storage cost (DLT-S)
- ACT-R target activation
- Left corner end of constituent (and length-weighted variants)
  ∘ +L, +L-WD, +L-DR, +L-DRV
- Left corner end of center embedding (+G)
- Left corner end of multiword center embedding (and length-weighted variants)
  ∘ +L+G, +L+G-WD, +L+J-DR, +L+J-DRV
- Left corner embedding depth
  ∘ Embedding depth, start of embedding

Families are selected for generalization-based testing if they contain at least one member (1) whose effect estimate goes in the expected direction and (2) which is statistically significant following Bonferroni correction on the training set. For families with multiple such members, only the best variant (in terms of exploratory *r*_diff_) is selected for generalization-based testing. To perform the generalization tests, all predictors selected for generalization set evaluation are included as fixed effects in a model fitted to the training set, and all nested ablations of these predictors are also fitted to the same set. Fitted models are then used to generate predictions on the (unseen) evaluation set, using permutation testing to evaluate each ablative comparison in terms of out-of-sample *r*_diff_. This analyses pipeline is schematized in **Figure 2B**.

**Figure 2:**
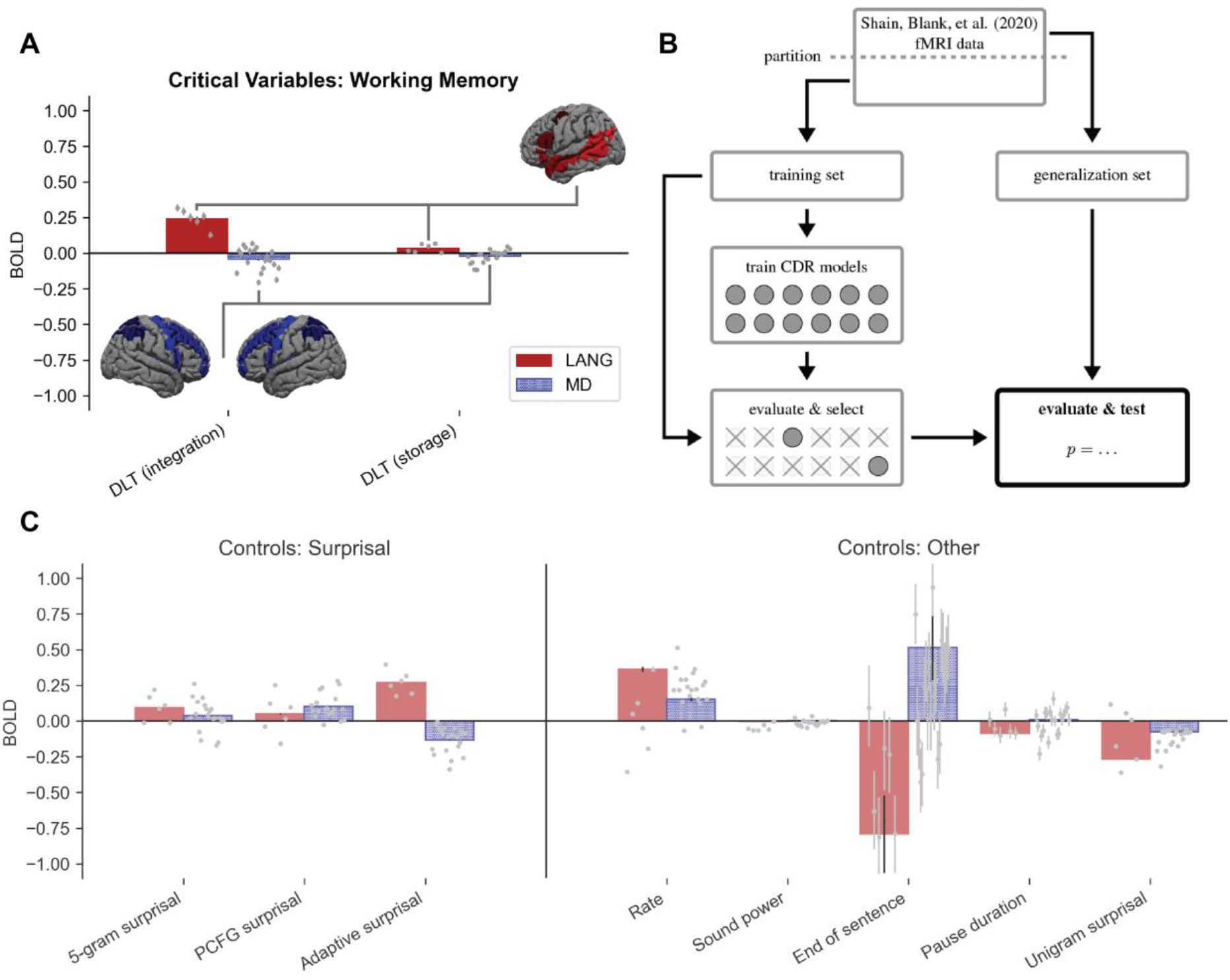
**A**. The critical working memory result, with reference estimates for surprisal variables and other controls shown in **C**. The LANG network shows a large positive estimate for DLT integration cost (DLT-VCM, comparable to or larger than the surprisal effect) and a weak positive estimate for DLT storage (DLT-S). The MD network’s estimates for both variables are weakly negative. fROIs individually replicate the critical DLT pattern and are plotted as points left-to-right in the following order: (LANG) LIFGorb, LIFG, LMFG, LAntTemp, LPostTemp, LAngG; (MD) LMFGorb, LMFG, LSFG, LIFGop, LPrecG, LmPFC, LInsula, LAntPar, LMidPar, LPostPar, RMFGorb, RMFG, RSFG, RIFGop, RPrecG, RmPFC, RInsula, RAntPar, RMidPar, RPostPar. (Note that estimates for the surprisal controls differ from those reported in Shain, Blank, et al. (2020). This is because models contain additional controls, especially *Adaptive Surprisal*, which overlaps with both of the other surprisal estimates and competes with them for variance. Surprisal effects are not tested because they are not relevant to our core claim.) Error bars show 95% Monte Carlo estimated variational Bayesian credible intervals. For reference, the group masks bounding the extent of the LANG and MD fROIs are shown projected onto the cortical surface. As explained in Methods, a small subset (10%) of voxels within each of these masks are selected in each participant based on the relevant localizer contrast. **B**. Schematic of analysis pipeline. The Shain, Blank, et al. (2020) fMRI data is partitioned into training and generalization sets. The training set is used to train multiple CDR models, two for each of the memory variables explored in this study (a full model that contains the variable as a fixed effect and an ablated model that lacks it). Variables whose full model (a) contains estimates that go in the predicted direction and (b) significantly outperforms the ablated model on the training set are selected for the critical evaluation, which deploys the pre-trained models to predict unseen responses in the generalization set and statistically evaluates the contribution of the selected variable to generalization performance.

### Accessibility

Data used in these analyses, including regressors, are available on OSF: https://osf.io/ah429/. Regressors were generated using the ModelBlocks repository: https://github.com/modelblocks/modelblocks-release. Code for reproducing the CDR regression analyses is public: https://github.com/coryshain/cdr. Detailed reproduction instructions will be provided at these locations upon acceptance. These experiments were not pre-registered.

## Results

### Exploratory Phase: Do WM Predictors Explain LANG or MD Network Activity in the Training Set?

Effect estimates and in-sample significance tests from the exploratory analysis of each functional network are given in **Table 1** (LANG) and **Table 2** (MD), with groupings of predictors into equivalent “families” shown by horizontal lines.

**Table 1:**
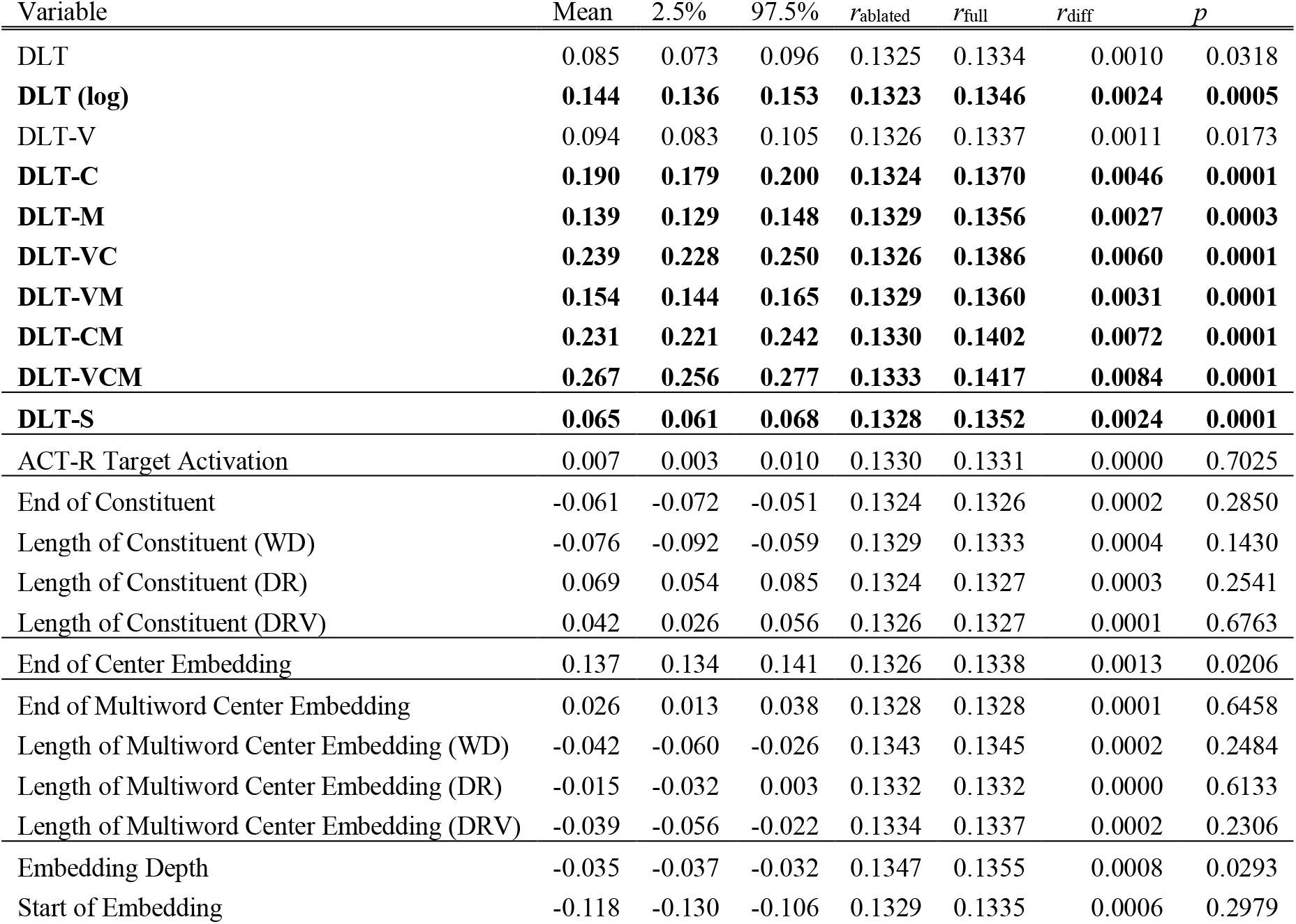
LANG exploratory results. V, C, and M suffixes denote the use of verb, coordination, and/or preceding modifier modifications to the original definition of DLT integration cost; DLT-S is storage cost. Effect estimates with 95% credible intervals, correlation levels of full and ablated models on the training set, and significance by paired permutation test of the improvement in training set correlation. Families of predictors are delineated by black horizontal lines. Variables that have the expected sign and are significant under 22-way Bonferroni correction are shown in bold (note that the expected sign of ACT-R Target Activation is negative, since processing costs should be lower for more activated targets). *r*_diff_ is the difference in Pearson correlation between true and predicted responses from a model containing a fixed effect for the linguistic variable (*r*_full_) to a model without one (*r*_ablated_).

**Table 2:**
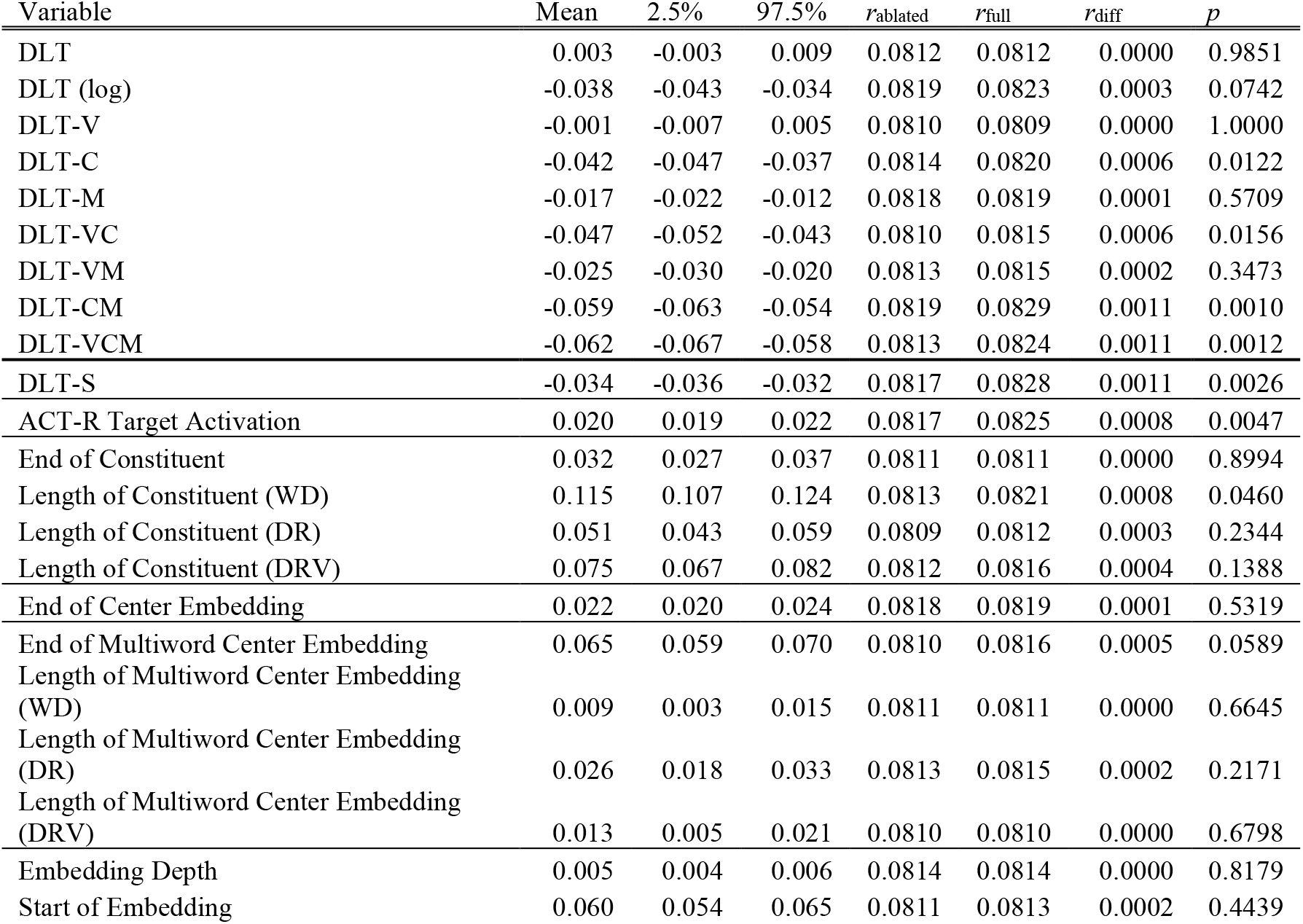
MD exploratory results. V, C, and M suffixes denote the use of verb, coordination, and/or preceding modifier modifications to the original definition of DLT integration cost; DLT-S is storage cost. Effect estimates with 95% credible intervals, correlation levels of full and ablated models on the training set, and significance by paired permutation test of the improvement in training set correlation. Families of predictors are delineated by horizontal lines. No variable both (1) has the expected sign (note that the expected sign of ACT-R Target Activation is negative, since processing costs should be lower for more activated targets) and (2) is significant under 22-way Bonferroni correction. *r*_diff_ is the difference in Pearson correlation between true and predicted responses from a model containing a fixed effect for the linguistic variable (*r*_full_) to a model without one (*r*_ablated_).

The DLT predictors are broadly descriptive of language network activity: 6/8 integration cost predictors and the storage cost predictor (DLT-S) yield both large significant increases in *r*_diff_ and comparatively large overall correlation with the true training response *r*_full_. The strongest variant of DLT integration cost is DLT-VCM. Although the log-transformed raw DLT predictor provides substantially stronger fit than DLT on its own, consistent with the hypothesis that the DLT overestimates the cost of long dependencies, it is still weaker than most of the other DLT variants, suggesting that these variants are not improving fit merely by discounting the cost of long dependencies. None of the other families of WM predictors is clearly associated with language network activity.

No predictor is significant in MD with the expected sign. Two variants of DLT integration cost (DLT-M and DLT-VCM) are significant but have a negative sign, indicating that BOLD signal in MD *decreases* proportionally to integration difficulty. This outcome is not consistent with the empirical predictions of the hypothesis that MD supports WM for language comprehension, though it is possibly instead consistent with “vascular steal” (Lee et al., 1995; Harel et al., 2002) and/or inhibition (Shmuel et al., 2006) driven by WM load in other brain regions (e.g. LANG).

In follow-up analyses, we addressed possible influences of using the flipped language localizer contrast (*nonwords > sentences*) to define the MD network by instead localizing MD using a *hard > easy* contrast in a spatial WM task (Fedorenko et al., 2013). Results were unchanged: no predictor has a significant effect in the expected direction (see OSF for details: https://osf.io/ah429/).

These exploratory results have several implications. First, they support the existence of syntactically-related WM load in the language network during naturalistic sentence comprehension and indicate that the DLT captures this WM signature better than more recent algorithmic-level models of WM in language (Lewis & Vasishth, 2005; Rasmussen & Schuler, 2018; Dotlačil, 2021). Second, they present a serious challenge to the hypothesis that the MD network – the most likely domain-general WM resource – is recruited for language-related WM operations: despite casting a broad net over theoretically motivated WM measures and performing the testing in-sample, the MD network does not show systematic correlates of WM demand. Based on these results, DLT-VCM and DLT-S are selected for evaluation on the (held-out) generalization set, to ensure that the reported patterns generalize. Models containing/ablating both predictors are fitted to the training set, and their contribution to *r* measures in the generalization set is used for evaluation and significance testing. Because the MD network does not register any clear signatures of WM effects, further generalization-based testing is unwarranted. MD models with the same structure as the full LANG network models are fitted simply in order to provide a direct comparison between estimates in the LANG vs. MD network.

These results also indicate that implementation variants in models of WM critically influence alignment with measures of human language processing load: only certain variants of the DLT register a strong effect, whereas the DLT as originally formulated and two other WM theories are not strongly associated with activity in either network. This outcome is important because, although the DLT does not commit to a parsing algorithm (Gibson, 2000), algorithmic-level theories like ACT-R and left corner parsing make empirical predictions that are in aggregate similar to those of the DLT (Lewis & Vasishth, 2005) and yet provide a poorer description of human neuronal timecourses. This raises three key questions for future research. (1) Are the gains from DLT integration cost and its variants significant over other theoretical models of WM in sentence processing? If so, (2) which aspects of the DLT (e.g., linear effects of dependency locality, a privileged status for nouns and verbs, etc.) give rise to those gains, and (3) how might the critical constructs be incorporated into existing algorithmic level sentence processing models to enable them to capture those gains?

### Generalization Phase

#### Do WM Predictors Explain Neural Activity in the Generalization Set?

Effect sizes (HRF integrals) by predictor in each network from the full model are plotted in **Figure 2A**. As shown, the DLT-VCM effect is strongly positive and the DLT-S effect is weakly positive in LANG, but both effects are slightly negative in MD.

The critical generalization (out-of-sample) analyses of DLT effects in the language network are given in **Table 4**. As shown, the integration cost predictor (DLT-VCM) contributes significantly to generalization *r*_diff_, both on its own and over the storage cost predictor (DLT-S). Storage cost effects are significant in isolation (DLT-S is significant over “neither”) but fail to improve statistically upon integration cost (DLT-S is not significant over DLT-VCM). Generalization-based tests of interactions of each of these predictors with *network* (a test of whether the LANG network exhibits a larger effect than the MD network) are significant for all comparisons, supporting a larger effect of each variable in the LANG network. In summary, the evidence from this study for DLT integration cost is strong, whereas the evidence for DLT storage cost is weaker (storage cost estimates are (a) positive, (b) significant by in-sample test, and (c) significantly larger by generalization-based tests than storage costs in MD; however, they do not pass the critical generalization-based test of difference from 0 in the presence of integration costs, and the existence of distinct storage costs is therefore not clearly supported by these results).

**Table 4:** Critical comparison. *p* values that are significant under 8-way Bonferroni correction (because 8 comparisons are tested) are shown in **bold**. For the LANG network (LANG (*p*) column), integration cost (DLT-VCM) significantly improves network generalization *r*_diff_ both alone and over storage cost (DLT-S), whereas DLT-S only contributes significantly to generalization *r*_diff_ in the absence of the DLT-VCM predictor (significant over “neither” but not over DLT-VCM). For the combined models (Interaction with *network* (*p*) column), the interaction of each variable with *network* significantly contributes to generalization *r*_diff_ in all comparisons, supporting a significantly larger effect of both variables in the language network than in the MD network.

|  | LANG (*p*) | Interaction with *network* (*p*) |
| --- | --- | --- |
| DLT-VCM over neither | **0.0001\*\*\*** | **0.0001\*\*\*** |
| DLT-S over neither | **0.0033\*\*\*** | **0.0013\*\*** |
| DLT-VCM over DLT-S | **0.0001\*\*\*** | **0.0001\*\*\*** |
| DLT-S over DLT-VCM | 0.3301 | **0.0007\*\*\*** |

Thus, whatever storage costs may exist, they appear to be considerably fainter than integration costs (smaller effects, weaker and less consistent improvements to fit), and a higher-powered study may be needed to tease the two types of cost apart convincingly (or to reject the distinction). In this way, our study reflects a fairly mixed literature on storage costs in sentence processing, with some studies reporting effects (King & Kutas, 1995; Fiebach et al., 2002; Chen et al., 2005; Ristic et al., 2021) and others failing to find any (Hakes et al., 1976; Van Dyke & Lewis, 2003).

#### How Well Do Models Perform Relative to Ceiling?

**Table 5** shows correlations between model predictions and true responses by network in both halves of the partition (training and evaluation), relative to a “ceiling” estimate of stimulus-driven correlation, computed as the correlation between (a) the responses in each region of each participant at each story exposure and (b) the average response of all other participants in that region, for that story. Consistent with prior studies (Blank & Fedorenko, 2017), the LANG network exhibits stronger language-driven synchronization across participants than the MD network (higher ceiling correlation). Our models also explain a greater share of that correlation in LANG vs. MD, especially on the out-of-sample evaluation set (39% relative correlation for LANG vs. 3% relative anticorrelation for MD).

**Table 5:** Correlation *r* of full model predictions with the true response compared to a “ceiling” measure correlating the true response with the mean response of all other participants for a particular story/fROI. “*r*-absolute” columns show absolute percent variance explained, while “*r*-relative” columns show the ratio of *r*-absolute to the ceiling.

|  | LANG |  | MD |  | Combined |  |
| --- | --- | --- | --- | --- | --- | --- |
|  | *r*-absolute | *r*-relative | *r*-absolute | *r*-relative | *r*-absolute | *r*-relative |
| Ceiling | 0.221 | 1.0 | 0.116 | 1.0 | 0.152 | 1.0 |
| Model (train) | 0.143 | 0.647 | 0.085 | 0.733 | 0.083 | 0.546 |
| Model (evaluation) | 0.086 | 0.389 | -0.003 | -0.026 | 0.048 | 0.316 |

#### Are WM Effects Localized to a Hub (or Hubs) within the LANG network?

Estimates also show a spatially distributed positive effect of DLT integration cost across the regions of the LANG network (gray points in **Figure 2A**) that systematically improves generalization quality (**Table 6**), indicating that all LANG fROIs are implicated to some extent in the processing costs associated with the DLT. Storage cost effects are less clear: although numerically positive DLT-S estimates are found in all language regions (**Figure 2A**), they do not systematically improve generalization quality (**Table 6**). No such pattern holds in MD: regional effects of both DLT variables cluster around 0 (gray points in **Figure 2A**), and the sign of *r*_diff_ across regions is roughly at chance. These results converge to support strong, spatially distributed sensitivity in the language-selective network to WM retrieval difficulty.

**Table 6:** Unique contributions of fixed effects for each of integration cost (DLT-VCM) and storage cost (DLT-S) to out-of-sample correlation improvement *r*_diff_ by language fROI (relative to “ceiling” performance in LANG of 0.221). DLT-VCM improves correlation with the true response in all (6/6) LANG regions, but DLT-S improves correlation with the true response in only 3/6 LANG regions.

| fROI | Network | DLT-VCM $r_{\text{diff}}\text{-relative}$ | DLT-S $r_{\text{diff}}\text{-relative}$ |
| --- | --- | --- | --- |
| IFGorb | LANG | 0.038 | 0.003 |
| IFG | LANG | 0.027 | -0.003 |
| MFG | LANG | 0.019 | -0.003 |
| AntTemp | LANG | 0.062 | 0.006 |
| PostTemp | LANG | 0.027 | 0.006 |
| AngG | LANG | 0.002 | -0.001 |

#### Are WM Effects Driven by Item-Level Confounds?

Because the data partition of Shain, Blank, et al. (2020) distributes materials across the training and evaluation sets, it is possible that item-level confounds may have affected our results in ways that generalize to the test set. To address this possibility, in a follow-up analysis we repartition the data so that the training and generalization sets are roughly equal in size but contain non-overlapping materials, and we re-run the critical analyses above. The result is unchanged: DLT-VCM and DLT-S estimates are positive with similar magnitudes to those reported in **Figure 2A**, DLT-VCM contributes significantly both on its own (p < 0.001) and in the presence of DLT-S (p < 0.001), and DLT-S contributes significantly in isolation (p < 0.004) but fails to contribute over DLT-VCM, even numerically (p = 1.0). The evidence from both analyses is consistent: WM retrieval difficulty registers in the language network, with little effect in the multiple-demand network. Effect estimates from this reanalysis are consistent with those in **Figure 2** and are available on OSF: https://osf.io/ah429/.

## Discussion

A standard view holds that typical human language processing involves (a) rich word-by-word structure building (b) via computationally-intensive operations in working memory (WM) (c) implemented in the same resources that support WM across domains. This study is motivated by prior challenges to the three core components of this view, namely (a) that the default mode of human language processing may be mostly approximated and shallow, especially in naturalistic settings (e.g., Frank & Bod, 2011), (b) that the main driver of language processing costs may be surprisal rather than WM demand (e.g., Levy, 2008), and (c) that the language system may rely on specialized (rather than domain-general) WM resources (e.g., Caplan & Waters, 1999). To investigate these hypotheses, we analyzed a large publicly available dataset of fMRI responses to naturalistic stories (Shain, Blank, et al., 2020) with respect to diverse theory-driven estimates of syntactically modulated WM demand (Gibson, 2000; Lewis & Vasishth, 2005; Rasmussen & Schuler, 2018) under rigorous controls for word predictability (Heafield et al., 2013; van Schijndel et al., 2013; van Schijndel & Linzen, 2018). To probe the nature of the computations in question, we examined neural responses in two functionally localized brain networks: the domain-specific, language-selective network (LANG; Fedorenko et al., 2010) and the domain-general, multiple-demand network (MD; Duncan, 2010), implicated in executive functions, including working memory.

Exploratory analyses of general theories of WM load in sentence comprehension, which posit *storage costs* associated with maintaining representations in WM and/or *integration costs* associated with retrieving representations from WM, identified clear effects in the LANG network of integration cost and weaker effects of storage costs over rigorous surprisal controls. No WM measures reliably characterized responses in the MD network.

Generalization tests on held-out data support both (1) integration cost (but not storage cost) as a generalizable correlate of activity in the LANG network, and (2) systematically larger effects of integration and storage costs in the LANG network than in the MD network. Further, integration costs are found across the different regions of the language network, supporting a broadly distributed WM system for language comprehension, rather than a spatially restricted “hub” or a set of hubs.

This pattern of results supports two broad inferences about the neural implementation of human sentence processing. ***First***, our results support the widely-held view that a core operation in human sentence processing is to encode and retrieve items in WM as required by the syntactic structure of sentences (Gibson, 2000; McElree et al., 2003; Lewis & Vasishth, 2005; Van Dyke & McElree, 2006; Rasmussen & Schuler, 2018), even in a naturalistic setting where behavioral evidence for such effects has been mixed in the presence of surprisal controls (Demberg & Keller, 2008; van Schijndel & Schuler, 2013; Shain & Schuler, 2018). And ***second***, our results challenge prior arguments that the WM operations supporting language comprehension draw on domain-general WM resources (Stowe et al., 1998; Fedorenko et al., 2006, 2007; Amici et al., 2007). Activity in the MD network—the most plausible candidate for implementing domain-general WM computations (Goldman-Rakic, 1988; Owen et al., 1990; Kimberg & Farah, 1993; Duncan & Owen, 2000; Prabhakaran et al., 2000; Cole & Schneider, 2007; Duncan, 2010; Gläscher et al., 2010; Rottschy et al., 2012; Camilleri et al., 2018; Assem et al., 2020)—shows no association with any of the WM measures explored here and shows significantly weaker associations with critical WM predictors than does the LANG network. Our results thus support the hypothesis that the WM operations required for language comprehension are be carried out by the brain regions that store linguistic knowledge (Caplan & Waters, 1999; Fiebach et al., 2001; Fedorenko & Shain, to appear).

The estimates of WM demand considered here are determined by the syntactic structure of sentences (as are all extant theories of WM demand in language comprehension to our knowledge). Therefore, the finding that integration cost robustly registers in the brain response to naturalistic stories provides indirect support for syntactic processing as a core subroutine of typical language comprehension. This result challenges to some extent prior arguments that representations computed during typical language comprehension are more approximate and shallow than is often assumed (Swets et al., 2008; Frank & Bod, 2011; Frank et al., 2015; Frank & Christiansen, 2018), although further research is needed to determine more precisely the level of syntactic detail present in human mental representations (e.g., Brennan, 2016; Brennan & Hale, 2019; Lopopolo et al., 2020). Our results also suggest a spatially distributed burden of syntactic processing across the regions of the language network (Bates et al., 1995; Caplan et al., 1996; Snider & Arnon, 2012; Toneva & Wehbe, 2019; Shain, Blank, et al., 2020; see Fedorenko et al., 2020 for review) and are not consistent with prior arguments for the existence of one or two dedicated syntactic processing centers (Vandenberghe et al., 2002; Hagoort, 2005; Friederici et al., 2006; Bemis & Pylkkänen, 2011; Pallier et al., 2011; Brennan et al., 2012; Matchin et al., 2017; Matchin & Hickok, 2020).

The WM effects shown here furthermore fail to be accounted for by multiple strong measures of word predictability, which has repeatedly been shown in prior work to describe naturalistic human sentence processing responses across modalities, including behavioral (Demberg & Keller, 2008; Frank & Bod, 2011; Fossum & Levy, 2012; Smith & Levy, 2013; van Schijndel & Schuler, 2015; Aurnhammer & Frank, 2019; Shain, 2019), electrophysiological (Frank et al., 2015; Armeni et al., 2019), and fMRI (Brennan et al., 2016; Henderson et al., 2016; Lopopolo et al., 2017; Shain, Blank, et al., 2020; Willems et al., 2015). In its strong form, surprisal theory (Hale, 2001; Levy, 2008) equates sentence comprehension with allocating activation between (potentially infinite) possible interpretations of the unfolding sentence, in proportion to their probability given the currently observed string. Under such a view, structured representations are assumed to be available, and the primary work of comprehension is (probabilistically) selecting among them. However, according to integration-based theories (Gibson, 2000; Lewis & Vasishth, 2005) incremental effort is required to *compute* the available interpretations in the first place (i.e. by storing, retrieving, and updating representations in memory). By showing integration costs that are not well explained by word predictability, our study joins recent arguments in favor of complementary roles played by integration and prediction in language comprehension (Levy et al., 2013; Ferreira & Chantavarin, 2018). Strong word predictability controls are of course a perpetually moving target: we cannot rule out the possibility that some other current or future statistical language model might explain apparent WM effects. However, such an objection effectively renders surprisal theory unfalsifiable. We have attempted to address such concerns by deploying a surprisal control (*adaptive surprisal*) that, at the time of writing, is, by psycholinguistic standards, recent, high performance, and cognitively motivated.

Notwithstanding, one recent variant of surprisal theory might offer an alternative explanation for our finding: *lossy context surprisal* (Futrell et al., 2021). Lossy context surprisal derives what we have termed “integration costs” as predictability effects by positing a memory store subject to a *progressive noise function*, whereby words are more likely to be forgotten the longer ago they occurred. Because dependencies can make words more predictable, forgetting renders words less predictable on average (and thus harder to process) when they terminate longer dependencies. Under this view, the driver of integration costs is not WM retrieval difficulty but rather prediction difficulty when critical prediction cues have been forgotten. Although lossy context surprisal places the burden of processing primarily on prediction rather than structure-building in WM, it shares with existing memory-based theories an emphasis on the centrality of WM for language processing. Because lossy context surprisal currently lacks a broad-coverage implementation, we cannot directly test its predictions for our study, and we leave further investigation to future work (though we note that our RNN-based surprisal control approximates lossy context surprisal and does not explain the main result).

Finally, our study bears on the role of cortical WM but does not rule out a domain-general role for sub-cortical structures in WM during language processing, especially the hippocampus (Olton et al., 1979; Olson et al., 2006; Yoon et al., 2008; Leszczynski, 2011; Yonelinas, 2013). While the contribution of MD regions to (non-linguistic) WM is well established, the role of hippocampus in WM (as opposed to long-term episodic memory) is less so, with some prior studies supporting a hippocampal role in WM (e.g., Lipska et al., 2002; Axmacher et al., 2010; Öztekin et al., 2010; Yonelinas, 2013; Schapiro et al., 2014) and others not (e.g., Baddeley & Warrington, 1970; Nadel & MacDonald, 1980; Shrager et al., 2008; Baddeley et al., 2010, 2011; Jeneson et al., 2010). We leave to future research investigation of possible subcortical WM contributions to core operations in language processing.

In conclusion, our study supports the existence of a distributed but domain-specific working memory resource that plays a core role in language comprehension, with little recruitment of domain-general working memory resources housed within the multiple demand network.

## Acknowledgments

This research was supported by NIH award R00-HD-057522 (E.F.) and by National Science Foundation grant #1816891 (W.S.). All views expressed are those of the authors and do not necessarily reflect the views of the National Science Foundation. E.F. was additionally supported by NIH awards R01-DC-016607 and R01-DC-016950 and by a grant from the Simons Foundation via the Simons Center for the Social Brain at MIT. The authors would also like to acknowledge the Athinoula A. Martinos Imaging Center at the McGovern Institute for Brain Research at the Massachusetts Institute of Technology (MIT), and the support team (Steven Shannon and Atsushi Takahashi). The authors also thank Nancy Kanwisher and Ted Gibson for recording the stories, EvLab members for help with data collection (especially Zach Mineroff and Alex Paunov), and Jakub Dotlačil for generously providing ACT-R predictors to use in our analyses.

